# Characterizing neuronal and population responses to electrical stimulation in the human hippocampo-cortical network

**DOI:** 10.1101/2024.11.28.625915

**Authors:** Mircea van der Plas, Frederic Roux, Ramesh Chelvarajah, Vijay Sawlani, Bernhard Staresina, Maria Wimber, David T. Rollings, Simon Hanslmayr

## Abstract

Direct electrical stimulation (DES) can advance our understanding of the intricate dynamics of the human hippocampo-neocortical network, which underlies complex cognitive processes such as spatial cognition and memory. This knowledge can help Neurotechnology to more effectively interface with this network and improve its functions. Here, we investigated the effects of DES in seven epilepsy patients under medical supervision recording single neuron activity alongside local field potentials to investigate neural responses to single pulses at different levels of granularity. Our results demonstrate that (i) single neurons respond to local electrical stimulation with a stereotypical pattern of short-lived increased excitation, followed by sustained inhibition, (ii) that input into the hippocampus from neocortex takes ∼100 ms, and (iii) that output from the hippocampus to the neocortex is gated by theta phase. These results are vital to inform the optimal choice of parameters for future electrical stimulation studies targeting the human memory system.

## 1. Introduction

The potential of direct electrical brain stimulation as a tool for studying the memory system has become apparent, ever since Penfield and Perot first reported that cortical stimulation within the temporal lobe of tested patients could trigger the vivid re-experience of distinct memories ^1^. Subsequent research further established the medial temporal lobe (MTL), and particularly the hippocampus which plays a central role in integrating information from across the brain, as crucial for our ability to encode and recall episodic memories ^2–4^. This has made the area an obvious target for brain stimulation, in the context of researching the human memory system, as well as for clinical studies hoping to improve memory in pathological cases (for a review see Kucewicz et al., 2022), and even developing a neural prosthesis for memory ^6^.

However, progress has been limited because the evidence for direct electrical stimulation (DES) within the Hippocampus and its neighbouring areas, to improve memory is rather mixed. While some studies reported memory improvements in stimulated patients ^7–10^, other studies found that DES decreases memory performance ^11–14^. There are many possible reasons that could contribute to these divergent findings, ranging from differences in stimulation protocols, to different target selection procedures. However, this inability to pinpoint the exact mechanism is tied to the fact that, despite its wide use, our knowledge about the exact neurophysiological effects of direct stimulation in the human hippocampus is very scarce. Not only do many studies differ in the protocols they use, even identical stimulation protocols can lead to very different responses within an area ^15^. To further add to the complication, network responses depend on the local state at the time of stimulation. For instance, the hippocampal response to neocortical stimulation (i.e. input) is modulated by the hippocampal (i.e. receiving) theta phase^16^. However, whether hippocampal output to the neocortex is also modulated by hippocampal theta phase is not clear to date.

Animal studies recording from single neurons during electrical stimulation found that the typical response appears to be a short-lived increase in excitation, followed by sustained inhibition of varying lengths ^17,18^. The exact mechanism behind this pattern is still unknown. It has been hypothesized that this typical response arises due to architectural properties of connectivity between excitatory and inhibitory neurons in the neocortex ^19^. However, the human hippocampus as well as it’s organisation, differs from the that of the neocortex and the hippocampus in commonly used animal models^20–22^. Characterizing the effects of DES is further complicated by the fact that responses are not necessarily limited to the stimulated area but can travel vast distances throughout the brain. For example, thalamic stimulation has been observed to cause the afore mentioned stereotypical excitation-inhibition response pattern in the primary visual cortex ^19,23,24^.

Another useful approach to characterising neural responses to stimulation is by examining the frequency spectrum of the local field potential (LFP) or more general intracranial electroencephalographical (iEEG) responses^25,26^. Information in the frequency spectrum can be divided into two major components. The periodic component, which reflects oscillatory activity that carries information on how information is being coded, and the aperiodic component, which consists of broadband effects which follow Brownian-noise patterns^27,28^. Different frequencies of the periodic component have been associated with different functions^29^. Importantly, oscillatory activity in the alpha and theta band has been reported to modulate information transmission within and between the hippocampus and relevant neocortical areas^30^. The aperiodic component is frequently treated as noise but can still provide relevant information on underlying brain mechanisms. For example, the slope of aperiodic component has been linked to local excitation-inhibition balance ^31^, while the offset of the aperiodic component has been associated with ongoing firing rates ^32^.

The current study investigated the effects of direct stimulation in the human hippocampus and wider MTL, to allow for a more targeted use of future instances of direct stimulation in humans. The stimulation was accompanied by recordings of intracranial electroencephalographic (iEEG) activity throughout the MTL and single unit activity (SUA) in the hippocampus. Stimulation occurred systematically at various stimulation sites, enabling the characterization of the neural dynamics of direct stimulation within the hippocampus, it’s surrounding white matter tracts, and within the temporal cortex.

Based on the literature described above we hypothesized that direct electrical stimulation would lead to changes in local firing rate responses. Such a response would most likely manifest itself as the stereotypical excitation-inhibition response pattern demonstrated throughout different areas and species.

Moreover, we assumed that the stimulation response would propagate throughout the brain to distant connected areas. Specifically, we expected that stimulation in the hippocampus causes responses in the neocortex and vice versa. Such responses would allow us to gauge the timing of information transfer between the hippocampus and neocortex. We hypothesized that we would measure a lag of ∼100-200 ms as originally suggested by a study investigating hippocampo-cortical directional connectivity ^33^. Based on oscillatory frameworks of the hippocampus and neocortex, we expected this neocortical response to depend on the phase of theta oscillations, i.e. the dominant local frequency in the stimulated area^16^.

## 2. Results

Stimulation occurred systematically from the inside out across each Behnke-Fried micro-macro electrode. This setup allowed for simultaneous neural stimulation along macro contacts and data acquisition on microwires in the hippocampus and macro contacts located within the hippocampus and several areas in the neocortex (see Fig 1). For the sake of clarity recordings made in the vicinity of the stimulated area will be referred to as local, while data recorded from an area that was not adjacent to the stimulation will be referred to as distal (for further details see methods).

**Fig 1:**
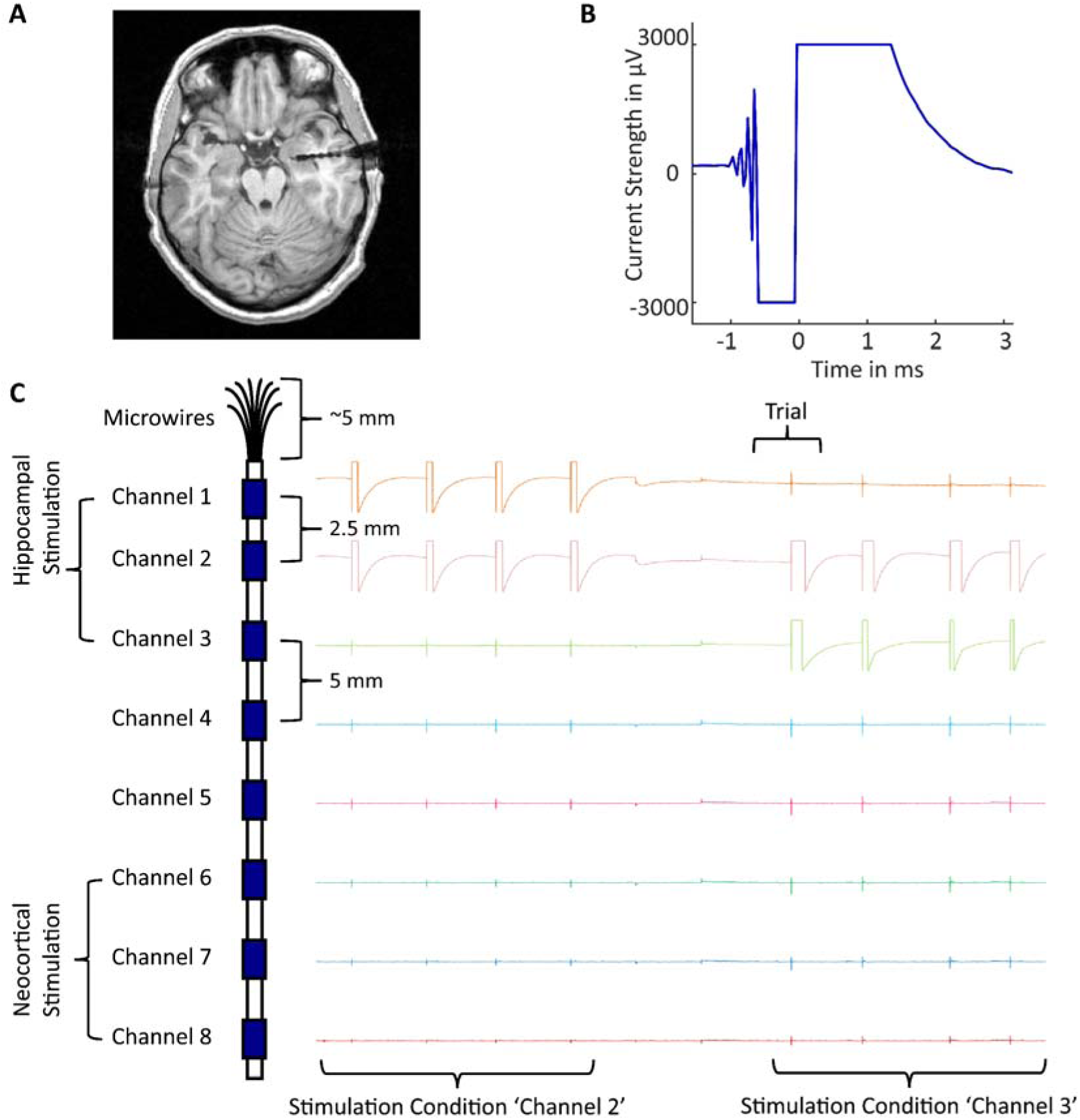
Illustration of the stimulation set-up. **A)** Example horizontal section of structural post-op MRI scan. The inserted electrode is visible as a black line with each ‘bead’ being a macro channel on the electrode. **B)** Example stimulation artifact as recorded on a micro-wire. The square, bimodal shape is clearly visible with the signal returning to baseline already after 3 ms. **C)** Diagram illustrating the Behnke-Fried electrode along with example signal from two different conditions before trial alignment (pictured time-window ∼90 seconds). The initial four stimulation pulses visible here were administered at the first and second channel, while the second set of pulses was administered between the second and third channel. Name for the condition is always derived from the outermost stimulated electrode of the pair.

### 2.1. Stereotypical SUA Response in the hippocampus

Of all detected single units (N = 136) across the seven patients, 29% showed a significant response to hippocampal stimulation (N= 39). A given unit was deemed responsive if its firing rate to local hippocampal stimulation fell outside the range of the 2.5^th^ and the 97.5^th^ percentile of the average baseline activity for time windows longer than 50 ms post stimulation (see Fig 2A for an example neuron). While this number of neurons is lower than initially expected it is still a highly significant number of neurons as determined by a binomial test (p < 0.0001). As hypothesized, the typical average response pattern consisted of a marked increase in firing rate (cluster size: 110 ms, p = 0.0017), followed by a prolonged silent period (cluster size: 141 ms, p > 0.0005) (see Fig 2B). This pattern was observed regardless of whether the microwires were located in epileptogenic sites or not (see supplementary figure 3 A and D for more information)

**Fig 2:**
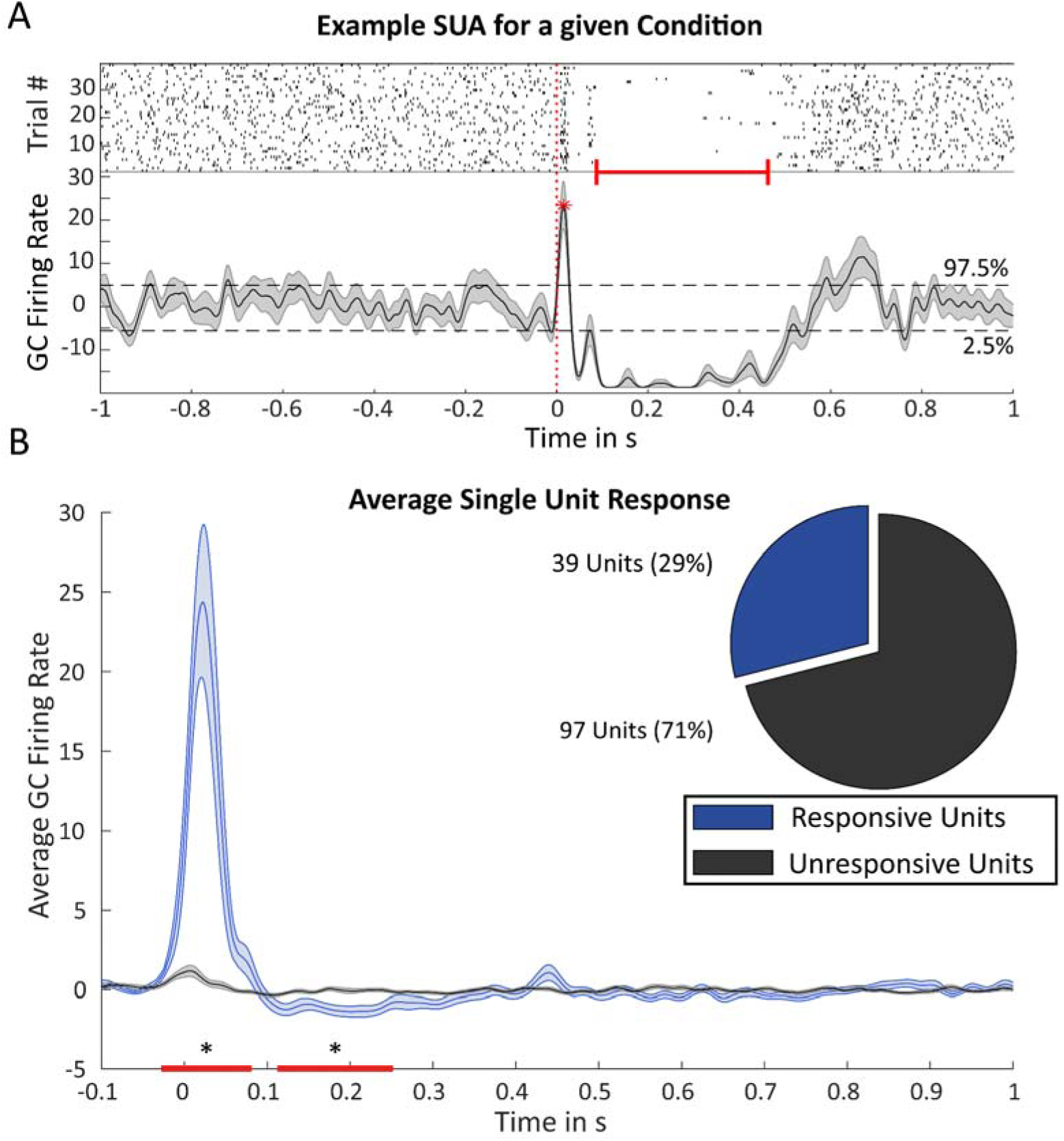
Average SUA response to SPES. **A)** Illustration of the procedure used to calculate the gaussian convolved (GC) firing rate per condition and the differentiation into responsive and unresponsive units. The vertical dotted red line indicates stimulation onset at time point 0. Top panel shows an example raster plot for a single unit during local stimulation in the hippocampus. The bottom panel shows the corresponding GC firing rate. Horizontally dashed black lines indicate the boundary percentiles as determined in the pre-stimulation period used for classifying responsive units. Red star indicates an example of a peak following stimulation. The red horizontal bar indicates the length of the silent period. Shaded area indicates standard deviation. **B)** GC firing rate across all conditions averaged separately for responsive units (black) and unresponsive units (blue). Red horizontal bar marked with an asterisk (*) indicates clusters of activity with a significant difference between the two groups as detected by a cluster-based permutation test comparing the responsive and unresponsive units. Shaded areas indicate standard error of the mean. Pie chart illustrates the ratio of responsiveness of all detected units.

Visual inspection of the raster plots of unit firing during hippocampal stimulation suggested that there might be a systematic pattern depending on the distance of stimulation to the respective unit (see Fig 3A for an example unit). To test whether this was a systematic pattern across all responsive units, we categorized them based on the Euclidian distance to the closest stimulated macro channel. For each condition the three following parameters were calculated: 1. The peak height, 2. the silent period length, 3. peak latency. Each of these three parameters was correlated against the calculated distance assigned to each condition after being normalized for each neuron. This analysis resulted in a significant negative correlation for peak height (r = −0.94, p => 0.0001) and log corrected silent duration (r = −0.93, p=> 0.0001) and peak latency (r = 0.79, p = 0.006) (all reported p-values are Bonferroni corrected; Fig 3B-C). These results suggest that with increasing distance peak height is reduced, and the silent period is shortened. Further inspection of the peak latency reveals that the negative correlation actually follows a step-function. The peak latency remains initially at the same value. For stimulation at distances greater than 15 mm, the latency sharply jumps forward to times around, and even preceding stimulus onset. Together with the data for the peak height and silent duration this suggests that most units do not respond to stimulation beyond 15 mm (the negative peak latencies are just noise detected by the peak detection algorithm in the absence of a peak resulting from stimulation). We conclude that the distance of stimulation does not affect the timing of the peak response. This implies that even at greater distances, the stimulation still directly influences the neurons, rather than through delays caused by synaptic transmission, which would have increased the peak latency at greater distances.

**Fig 3:**
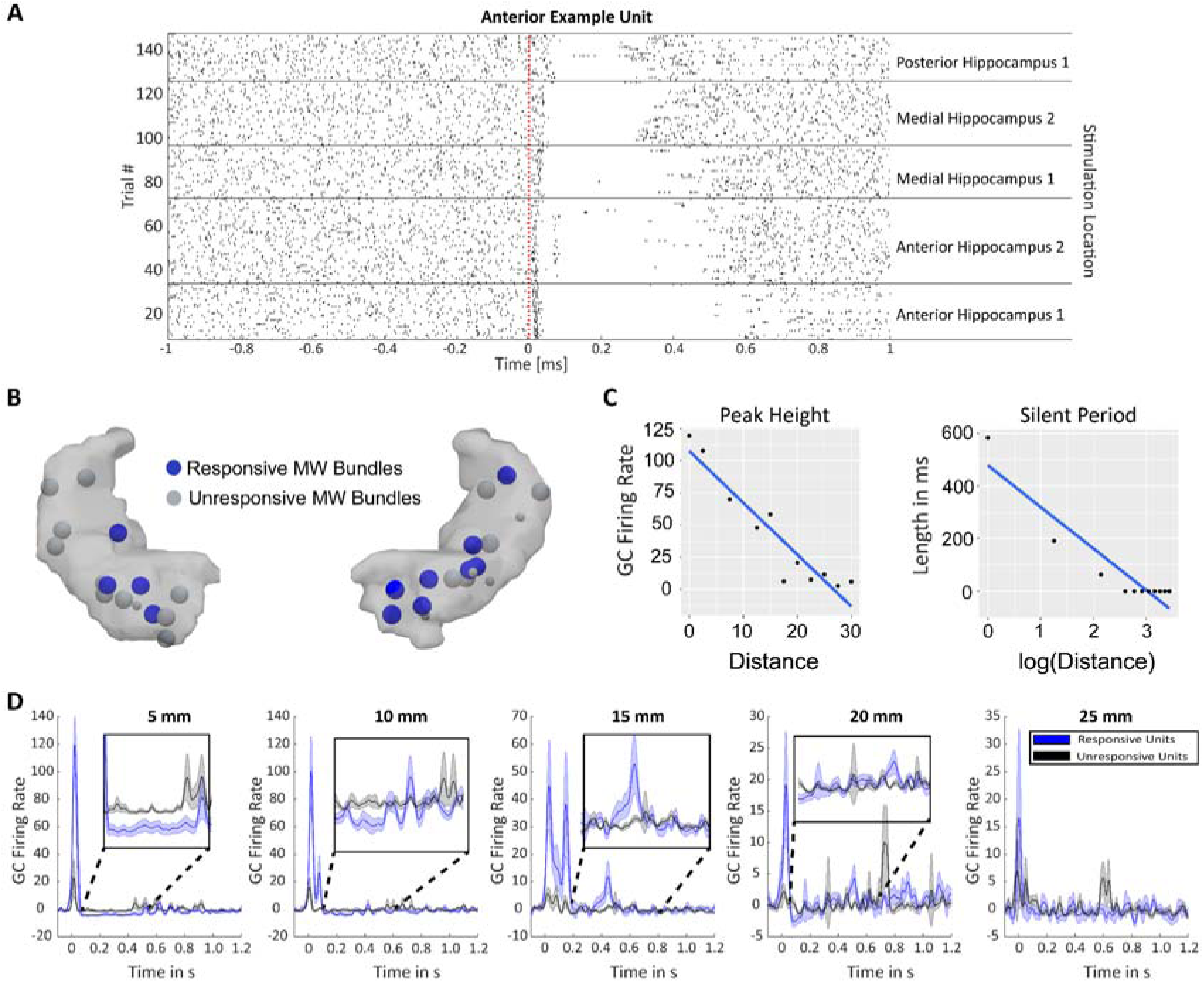
SUA response as dependent on distance of stimulation site. **A)** Firing rates of an example unit in the anterior hippocampus during stimulation at different hippocampal stimulation sites. Dotted red line indicates stimulation onset at time point 0. Labels on the right denote the stimulation site. **B)** Anatomical locations of the different hippocampally implanted micro-wires projected onto MNI space. Microwire bundles that exhibited a significant response to stimulation are highlighted in blue. Hippocampal electrodes that did not receive local stimulation are represented by a smaller circle. **C)** Scatter plots of each parameter of interest respectively: peak height and length of silent period. Each dot represents the parameter as extracted from the average GC firing rate for a condition at a given binned distance across units. **D)** GC firing rates plotted per binned distance of the recording wire to the stimulation channel separated by responsive and non-responsive neurons. Inset serves to highlight the parts of the post-stimulation response with a low amplitude. Shaded area indicates standard error of the mean.

To visualize these relationships averaged gaussian convolved (GC) firing rates were binned in 5 mm distance (from stimulation site) steps (see Fig 3D). These results confirm that the peak height and silent period length decrease with distance. Regarding peak latency, however, it appears that for distances exceeding 20 mm the peak in the responsive neurons does not exceed baseline level (i.e. non-responsive neurons). We also performed a control analysis to ensure that the observed activity does not result from direct propagation of the electric current conducted through the CSF as opposed to neuronal activity (see Supplementary Figure 2).

In summary, analysis of the SUA response reveals that a single pulse of DES elicits a response pattern consisting of an initial excitation, followed by a period of sustained inhibition. The intensity/length of both these responses is dependent on the distance to the stimulation site.

### 2.2. DES LFP response follows a stereotypical pattern

Next, we analysed the LFP responses recorded by the microwires. Since the activity on each Behnke Fried micro-wire bundle is highly correlated we grouped the responses to stimulation by bundle rather than individual microwire electrodes. As with the SUA analysis, we first determined the ratio of wires that showed a significant response to local hippocampal stimulation. For this, we selected only channels that showed a robust sustained response to local stimulation by comparing the maximum peak absolute values in a 400 ms post stimulation window, compared to a pre-stimulus window of equivalent length using a two-sided paired-samples t-test (see Methods section for details). We also verified through visual inspection that such a response occurred reliably on a single trial basis. Altogether, 44.44% (12/27) of the electrode bundles located in the hippocampus exhibited a response to local stimulation (see Figure 3B for the locations). Of the responsive bundles of wires, only one was located in posterior locations, with anterior and medial bundles showing more consistent responses to stimulation. Moreover, just like in the single-unit spike data, the pattern was not affected by location in epileptogenic tissue (see supplementary figure 3).

Having determined the response to local stimulation we looked at the ratio of electrodes exhibiting a response to distal stimulation (i.e. stimulation in the neocortex for hippocampal recording). To this end, we considered the microwires (bundles) and macro (iEEG) contacts separately. Concerning the microwires, merely 12.13% (4/32) of the hippocampal bundles showed a significant response to neocortical stimulation. In contrast, 45.83% (11/24) of the hippocampal macro channels showed a significant response to neocortical stimulation. Therefore, the macro channels were used to determine the signal delay between the medio-temporal neocortical areas and the hippocampus. To determine general response patterns, we averaged the ERPs based on local and distal hippocampal responses. The general LFP response in the hippocampus to local stimulation is characterized by an initial positive deflection which is followed by a longer negative deflection after ∼400 ms (Fig 4A). This pattern was also observed in hippocampal channels when stimulation occurred at distal, neocortical sites (Fig 4C). Interestingly, the peaks in the stimulation evoked potentials appeared at shorter latencies in local compared to distal stimulation. This likely reflects conduction delays of inputs from neocortical sites to the hippocampus.

**Fig 4:**
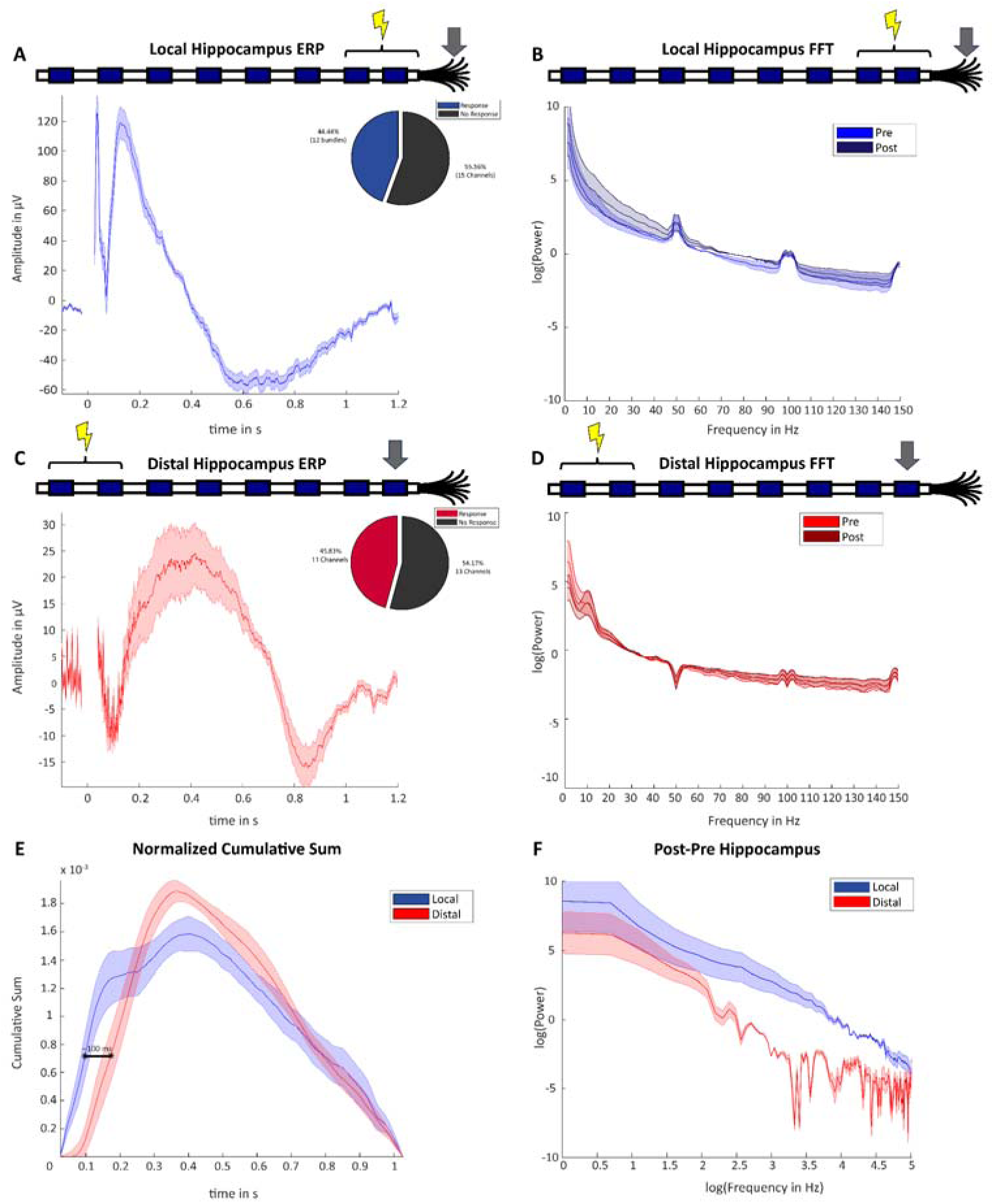
iEEG responses to Local and Distal stimulation. Shaded area indicates standard error of the mean. **A)** Average ERP responses over all local stimulation conditions for the hippocampus (MW data). **B)** Average fast fourier log-transformed (FFT) data in the local stimulation condition (MW data) for a 1 second pre stimulation and a 1 second post stimulation time-window **C)** Average ERP response over all distal stimulation conditions for the hippocampus (MC data). **D)** Average fast fourier log-transformed (FFT) data in the Distal stimulation condition (MW data) for a 1 second pre stimulation and a 1 second post stimulation time-window. **E)** Average slope of the normalized cumulative sum of the ERP responses for each condition. The black line serves to highlight the lag between ERP responses to local vs distal stimulation (∼100ms). **F)** Difference of the post-pre stimulus conditions for the local and distal stimulation conditions respectively.

**Fig 5:**
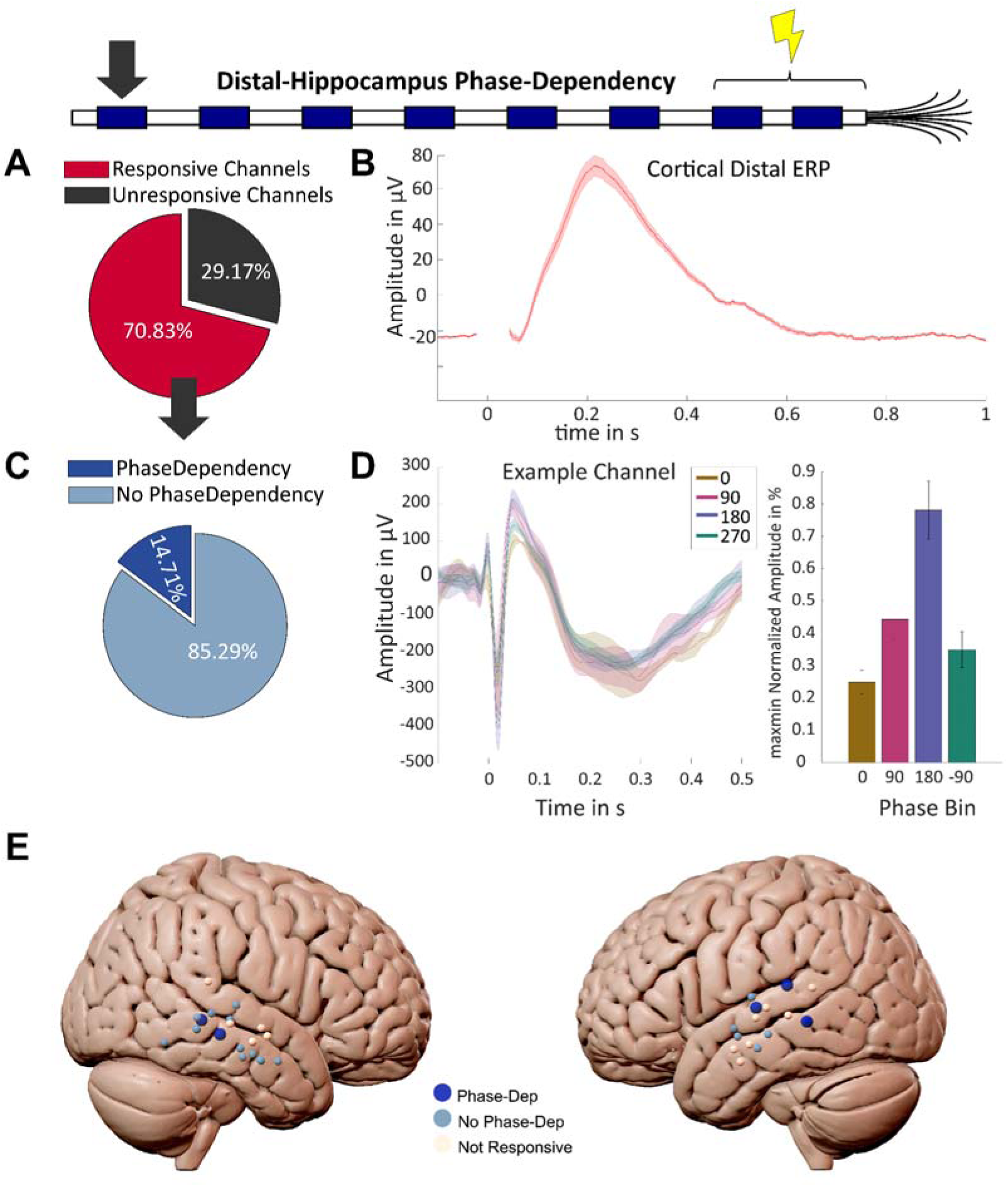
Distal Phase-Dependence Results. **A)** Proportion of Neocortical channels responding to hippocampal stimulation**. B)** ERP of the neocortical response to hippocampal stimulation **C)** Proportion of stimulation responsive lateral-temporal neocortical channels exhibiting stimulation phase dependency in response to hippocampal stimulation. **D)**. Example ERPs of a responsive channel, where each line represents an ERP binned according to the ongoing phase in the hippocampus. The colour indicates the corresponding hippocampal theta-phase. Histogram insert provides an alternative visualisation depicting the mean trial amplitude for each hippocampal theta phase bin, where each bin is centred around the phase indicated on the x-axis +-45 degrees **E)** Locations of the neocortical channels used in the phase-dependency analysis. Blue spheres indicate channels show a response to hippocampal stimulation, while white spheres indicate unresponsive channels. Stimulation response inducing channels are further subdivided into channels whose response depends on the ongoing hippocampal theta phase in the hippocampus (Dark blue) and channels that do not show this phase-dependency (light blue).

To further explore these delays in ERP responses, we computed the cumulative sum for each condition. To make sure that the cumulative sums are comparable across different types of electrodes, amplitudes and polarities, the cumulative sum was normalized to have an area of 1. Then, we calculated the slope of this normalised cumulative sum (see Methods), which results in a curve that represents the rate of change at any given time point. This approach allowed us to establish whether there was a systematic lag between the responses in local vs the distal stimulation conditions in an unbiased manner (Fig 4E). Thus, any differences in these slopes would relate exclusively to the timing and not the magnitude of the responses, which is necessary because trivially local stimulation results in higher responses than distal stimulation. A cluster-based permutation test comparing local vs distal cumulative sums of the responses revealed that the local ERP response develops faster than the distal ERP response in the hippocampus (p = 0.0247) by ∼100 ms for the first 200 ms.

For the frequency analysis the data was split into aperiodic and periodic components (for the log-transformed frequency spectra see Fig 4B, D, F). Due, to the lack of consistent oscillatory peaks in the data, no reliable statistical comparison in that regard was feasible. Moreover, disentangling the oscillatory activity from ongoing ERPs would complicate the interpretation of this data even more. The aperiodic components were analysed using a 2 (Stimulation: pre-vs post-stimulus) x 2 (Distance: local vs distal) ANOVA for the offset and exponent respectively (see Appendix 3 for the full output). For both the offset and exponent of the aperiodic activity a significant interaction effect was observed (F(1,20)=51.266 p<0.001 η^2^_p_=0.081; F(1,20)=50.755 p<0.001, η^2^_p_=0.176, respectively).

The direction of the effects suggest that stimulation leads to a decrease in the offset and a steepening of the spectrum through the increased exponent. These effects were much more pronounced in the local vs. the distal hippocampus condition. Whether the difference in magnitude of the effects is purely due to the distance of the stimulation or due to the properties of microwires vs sEEG electrode channels is, however, unclear. Therefore, these results should be interpreted with caution.

### 2.3. Phase-dependence of Hippocampal output to Neocortex

Investigation of the neocortical channels revealed that 70.83% (17/24) of the neocortical channels responded to hippocampal stimulation, which is considerably more compared to macro channels in the hippocampus responding to neocortical stimulation. But, are these distal effects modulated by the current state of the network, for instance the oscillatory phase at the stimulated area? Intuitively, it can be assumed that if stimulation is applied during states of high excitability, then it will propagate more strongly in the network compared to when it is applied during periods of low excitability^34–36^. Following previous work, we hypothesized theta oscillations, which is the dominant frequency in the human hippocampus, to play such a gating role^37,38^.

For this analysis we only included those channel pairs that showed a significant response to distal stimulation. When analysing neocortical responses to hippocampal stimulation 34 channel-pairs out of 65 (52.31%) showed a significant response (see methods for more information). Of this subset 14.71% exhibited evidence of a theta (4-8 Hz) phase dependency following hippocampal stimulation, which means that the ERP in the neocortex following hippocampal stimulation was modulated by the ongoing theta phase in the hippocampus (for an example phase-dependant neocortical response see Fig 4A-B). A follow-up binomial test suggested that this is a significantly larger proportion than expected by chance (p = 0.026). Note that this analysis uses ICA to extract the dominant theta component, and therefore its phase, from microwires in the hippocampus in an unbiased and data-driven manner (see Michelmann et al., 2018). However, one disadvantage of ICA is that it is non-deterministic therefore the exact values can change from run to run.

To verify that these results are specific to the theta band we performed two control analyses with a higher and a lower frequency of interest that do not overlap in phase with the original theta frequency by multiplying or dividing the theta frequency by the golden mean (1.618)^39^. Neither of these analyses revealed a significant proportion of phase dependency between the hippocampus and neocortex (p = 0.512 and p = 0.488 for frequencies above and below the originally tested theta frequency).

## 3. Discussion

In this study we used direct electrical stimulation alongside intracranial recordings to investigate (i) response patterns of hippocampal neurons and populations of neurons to DES, (ii) the time delay of neocortical input into the hippocampus (iii) whether hippocampal output to the neocortex is modulated by ongoing local oscillations. At the level of single neurons, we found that the typical response to DES in the hippocampus is characterized by a brief initial period of excitation followed by a prolonged period of inhibition. At the population level we were able to show that the conduction delay of neocortical input into the hippocampus is at a timescale of about ∼100 ms and that the output of the hippocampus to the neocortex is modulated by the phase of ongoing hippocampal theta oscillations.

The excitation-inhibition response pattern in the hippocampus in response to single pulse DES is in line with what has been observed in work with rodents and non-human primates^17–19^. However, the exact origin of the prolonged inhibitory silent period is unknown. One possibility is that DES initially mainly drives excitatory pyramidal neurons. This could be due to them being more numerous and larger compared to the smaller inhibitory interneurons which makes these neurons more receptive to the relatively large electrical gradients induced by the DES^40,41^. This would then lead to the observed short but strong firing rate increase in response to direct electrical stimulations. Such a burst of excitatory activity would drive the inhibitory neurons in the network to start a compensatory response which then inhibits neural firing, resulting in an overcorrection of the strong excitation with a long period of inhibition. In this previous interpretation the inhibition is a reaction to an initial burst of excitation. Another possible explanation for the observed pattern is that instead of a causal chain, the pattern arises from a general increase in neural firing across all neuron types at the same time. The pattern would then reflect the different firing properties/time-courses of excitatory pyramidal and inhibitory inter neurons. The initial excitation would be a result of the more numerous burst firing typical of pyramidal neuron drowning out the more sustained response of the interneurons, which subsides due to the prolonged interneuron activity that was triggered at the same time.

Regardless of the specifics, the above accounts on the underlying mechanism of the inhibitory response would further help explain why stimulation leads to a decreased offset of the aperiodic component as well as a steepening of the aperiodic component as evidenced by an increased exponent. Decreased offsets (i.e. broadband power decreases) have been associated with decreased spiking activity ^31,32^. While this result makes intuitive sense with the other results presented in this paper, we would like to highlight the caveat that we cannot rule out that the differences across the frequency spectra are not due to inherent properties of the micro-wires vs sEEG channels. These findings should thus be interpreted with caution.

Another interesting finding is that the inhibitory effect resulting from stimulation is retained across large distances; Stimulation in the hippocampus evoked similar (albeit smaller) response patterns in the neocortex as it did for local stimulation. This is consistent with findings such as by Logothetis et al. ^19^, where thalamic stimulation produced inhibitory responses in occipital visual regions. This property of producing distal responses was leveraged in this study to obtain an estimate of the conduction delay from the neocortex to the hippocampus, i.e. neocortical input into the hippocampus. The average delays observed in the cumulative sum analysis suggest a delay of ∼100 ms.

Estimating such delays of information transmission between the hippocampus and neocortex helps to narrow down the parameter space in the context of human episodic memory. Most models of memory involve some sort of interaction between the hippocampus and the neocortex ^30,42,43^. One could use single pulses as a model for information transmission that occurs naturally during memory processes. For example, a pulse evoking a response in the hippocampus could be analogous to the information transmission that occurs during encoding of memories. Here too, the neocortical areas process the information first before sending this signal to the hippocampus which binds, and possibly stores the information from all across the cortex somehow.

The ∼100 ms transduction delays for hippocampal activity, measured here correspond to neocortical and hippocampal activity delays during memory encoding as measured in a in a study conducted by Griffiths et al. ^33^. In this study they were also able to measure delays in the other directions, i.e. from the hippocampus to the neocortex or hippocampal output. This output from the hippocampus to the neocortex took between 200-300 ms which suggests that the input into the hippocampus from the neocortex takes approximately half the time as the output from the hippocampus to the neocortex. If one equates input with memory encoding, and output with memory retrieval, it would suggest that the delays observed during encoding approximately equate the time of signal transmission from the neocortex to the hippocampus, while retrieval requires extra processing steps leading to longer delays of hippocampal output to neocortex. Pattern completion, which is assumed to happen in recursive loops in area CA3, could be one candidate mechanism for lengthening this processing time^44,45^.

Lastly, we found that hippocampal theta phase modulates the stimulation response in the neocortex, i.e. the hippocampal output into the neocortex. Hippocampal theta oscillations have been suggested to regulate memory by gating information transfer to and from the hippocampus ^46,47^. Specifically, it has been hypothesized that certain phases bias an area to receive input, while other phases bias an area to transmit information and bias other areas to be receptive to that information. Our result is in line with these frameworks. More specifically, our results are consistent with the Separate Phase of Encoding and Recall (SPEAR) model put forward by Hasselmo (2005) which suggests that hippocampal output is modulated by theta phase. Going beyond basic science, this result is also important for Neurotechnology, because it shows that in order for DES to effectively modulate network activity, it has to be delivered at the right phase. This could be delivered by ‘read-and-write’ devices which deliver phase-dependent stimulation.

This result builds on a previous study by Lurie et al. ^16^, who showed that distal hippocampal responses are modulated by ongoing theta phase during neo-cortical stimulation in the human brain, and extend these findings in the following ways. First of all, we combine microwire with macro-channel data which allows to record single neuron responses during stimulation and also to separate the artefact induced by the stimulation from neural activity because the stimulation artifact is very short-lived in the microwire recordings. Second, we show that theta phase in the hippocampus modulates the output to neocortical areas. One difference from Lurie et al is that our study shows weaker responses in the hippocampus to neocortical stimulation. Possible reasons for this could be the much higher impedance of microwires which means that they are picking up a much more localised signal in the hippocampus than macro channels (as used in Lurie et al). This could also be the reason for why we do not see the phase-dependence resulting from cortical stimulation in the hippocampus, as found in Lurie et al.

Another difference between this study and Lurie et al. is their use of a low-frequency repetitive stimulation paradigm of 0.5 and 1 Hz stimulation, whereas we used a single pulse stimulation where stimuli occurred non-rhythmically at long time intervals (between 5 and 10 seconds). The fact that phase-dependent activity can be detected across the hippocampus and neocortex using different stimulation paradigms reinforces the importance of considering ongoing oscillations during stimulation in general. This is further strengthened by the fact that this relationship was determined in this study by determining the individual theta frequency and phase on a trial-by-trial basis.

Together, our results add to our understanding of the hippocampo-neocortical network dynamics in characterising how single neurons in the hippocampus respond to local stimulation, mapping the input latency from the neocortex to the hippocampus and showing that hippocampal output is gated by theta phase. These results are important because they advance inform computational models of human hippocampo-neocortical networks underlying memory and spatial cognition. They are also important for Neurotechnology to inform parameter choices of stimulation devices that aim to improve memory via electrical stimulation.

## 4. Methods (Online)

### 4.1. Participants

The data was acquired from seven patients (60% female; age: 35+-10.8; range: 23 to 53 y) under medical observation due to pharmacologically intractable epilepsy at the Queen Elizabeth Hospital Birmingham (QEHB) over the course of a total of twelve sessions. Patients were implanted with 2-6 Behnke-Fried hybrid micro-macro electrodes (Ad-Tech Medical Instrument Corporation, Oak Creek, WI) per person terminating in the hippocampus for a period of 10-14 days to evaluate the possibility for a surgical resection as treatment ^48^. The location of the electrodes was based entirely on clinical requirements. Electrode locations were confirmed visually using pre- and post-op structural magnetic resonance imaging (MRI) scans (For an example of an electrode as registered by an MRI scan, see Fig 1 A). Written informed consent was obtained in accordance with the Declaration of Helsinki.

### 4.2. Data recording and pre-processing

#### 4.2.1. iEEG

Each Behnke-Fried electrode consisted of eight macro-contacts and one microwire electrode bundle from which the iEEG signal was recorded (see Fig 1 C for a diagram). The microwire bundles comprised 8 recording wires and a ninth reference wire. The microwire bundles recorded hippocampal SUA and local field potential (LFP) at a 32 kHz sampling rate through a 64-channel ATLAS Neurophysiology system (Neuralynx Inc.).

The iEEG data acquired by the macro-contacts was recorded at a 1024 Hz sampling rate. Most iEEG pre-processing steps were performed in MATLAB, using the Fieldtrip toolbox, unless otherwise specified ^49^.

After aligning the microwire iEEG/LFP data (MW) and the macro-contact iEEG data (MC), the MW data was downsampled to 1000 Hz. Line noise was partly removed from the data using the Chronux toolbox ^50^. All data was subsequently demeaned and cut into 6 second epochs (3 s pre-stimulation artifact 3 s post-stimulation artifact) and manually checked for artifacts resulting from noise or epileptic discharges. The period 20 ms before and 24 ms after stimulation onset was removed for all subsequent analyses based on simulations of the MW stimulation artifact showing that filter ringing persists for about 15 ms around the stimulation artifact. The trials were then sorted according to stimulation site as separate conditions. Each condition name is based on the lateral channel name (furthest away from the neocortex) at which stimulation was applied. The same steps were performed for the MC data, with the additional step of re-referencing all contacts per electrode to a channel located in white matter determined through visual inspection of the respective T1 weighted anatomical scans.

The conditions are split into local and distal conditions. Local refers to conditions where the recordings originate from the immediate vicinity of the source of stimulation (E.g. hippocampal MW activity during hippocampal stimulation). Distal stimulation on the other hand refers to recordings obtained from an area that is not within the same region of grey matter (E.g. hippocampal MW activity during stimulation in the neocortex).

For the calculation of event-related potentials (ERPs), iEEG activity was averaged per bundle (i.e. across the eight highly correlated micro-wirse) after removing noisy wires from the data. Activity from the bundles were subsequently averaged to create the grand-average ERPs. To make the ERPs of local and distal activity more comparable, the cumulative sum of the respective ERPs was calculated for the time-window running from stimulation onset to 1.2 s post stimulation.

The cumulative sum values were calculated over the ERP period. These values were normalized by dividing all values by the maximum cumulative sum value, creating an incrementally increasing value from 0 at 0.026 s to 1 at 1.026 s post stimulation. Thus, one could see the result as representing the percentage value of the completed ERP per time unit. To allow for comparisons across electrodes, the cumulative sum values were normalized to all add up to one by dividing the cumulative sum by it’s last maximum/last value. These cumulative sum values from different ERPs can be compared to each other for every given time point. If a cumulative sum value is higher relative to another, this is an indication this ERP has progressed faster through its total response than another. If it is lower, it is an indication that the ERP’s response lagging behind in time. This allowed the comparison of time-courses of the ERPs for local and distal stimulation independent of polarity and precise pattern of the signal^51^.

The frequency analysis was performed by using a multi-taper (specifically discrete prolate spheroidal sequences) fast Fourier transform with 1 Hz of spectral smoothing for frequencies ranging from 1 Hz to 150 Hz. The time window used was one second before and one second after the SPES pulse (pre: −1.022 - −0.022; post: 0.026:1.026). The resulting Frequency power-spectra were analysed further by separating the aperiodic and possible periodic components using the FOOOF toolbox ^27^.

#### 4.2.2. SUA

SUA was extracted by performing spike detection and assisted spike sorting using the Waveclus toolbox ^52^. The same trial definition used for the MW and MC data was applied to the SUA data. The SUA data for each unit was also separated by stimulation site (for an example see Appendix 1). The data per stimulation site was then binned to correspond to a 1000 Hz sampling rate and convolved with a Gaussian kernel yielding a Gaussian convolved (GC) firing rate for each condition (window size: 50 ms). The GC firing rate was normalized per unit by a z-transformation where each trial was subtracted with the average baseline firing rate pre-stimulus (−1000ms -> −250 ms relative to stimulation onset) over all trials for a given neuron and divided by the corresponding standard deviation derived over the baseline period of all trials for the normalized neuron.

Units were categorized as responsive or unresponsive to stimulation. A responsive unit was defined as a unit whose average GC firing rate to local hippocampal stimulation would exceed the 97.5^th^ percentile or would fall below the 2.5^th^ percentile of the average baseline activity after stimulation for at least 50 ms (see Fig 2A for an illustration). Responsive units were further analysed by subdividing the activity for each unit by the distance of stimulation. The estimated distance (d) between the unit and stimulation channel was defined was calculated as d = √(x2 - x1)^2^ + (y2 - y1)^2^ + (z2 - z1)^2^, where x1, y1, and z1 are the respective anatomical coordinates as derived from the anatomical localization for the stimulation site while x2,y2, and z2 are the respective estimated coordinates of the microwire from which the unit was recorded. This estimation was based on extrapolation of insertion angle of the electrode and the length of the cut microwires.

### 4.3. Stimulation

Single pulse electrical stimulation (SPES) was applied repeatedly between a pair of neighbouring macro-channels. Each trial contains a 1 ms biphasic pulse (current strength: 8 mA, charge density: 96 μC/cm2/pulse). A pulse was applied 10-15 times per pair (see Fig 1B for an example of the stimulation artifact as recorded in the microwires). Stimulation always started at the most medial channel pair (i.e. in the hippocampus) and moved laterally along the electrode. Stimulation always started at the most anterior electrode and ended at the most posterior electrode (for an illustration see Fig 1C). Due to signal saturation resulting from electrical stimulation, MC activity at the stimulating channels is not interpretable.

### 4.4. Statistical Analysis

To compare the activity of the responsive and non-responsive neurons and to find differences in the cumulative sum of the ERPs, cluster-based permutations with 10 000 permutations were performed ^53^. For the SUA activity the analysis was run as a two-tailed test with a cluster alpha of 0.05 on a time-window running from 0.1 s pre-stimulation to 1 s post-stimulation to be able to detect both increases and decreases in firing rate. The same analysis was performed for the cumulative sum, with the difference that the test was performed one-tailed as it was expected that the cumulative sum of local stimulation ERPs would always lead over the distal counterparts. Furthermore, only the post-stimulation time-window was tested from 0.026 to 1.026 seconds. The average difference in cumulative sum was calculated by averaging the time-difference within the resulting significant cluster.

For the Frequency analysis separate 2×2 (Stimulation: Local x Distal; Time: Pre x Post) Analyses of Variance (ANOVAs) were performed for the periodic and aperiodic components resulting from the FOOOF analysis. A Bonferroni corrected alpha was used for these ANOVAs.

### 4.5. Phase Dependency Analysis

For the phase dependency analysis we chose to perform an analysis based on a circular to linear correlation between the hippocampal phase values and the amplitude in the neocortex recorded by the macro-channels ^54^. To do this, first a subset of hippocampal and (temporal) neocortical channel pairs were selected. For a channel pair to be selected the neocortical channel had to show a significant response to hippocampal stimulation, which was operationalized by looking for an average absolute maximum value in a 400 ms window post-stimulation that was significantly higher than an equivalent value in a 400 ms window pre-stimulation as determined by a two-sided paired-samples t-test.

Once the responsive channels were paired with a given stimulation site in the hippocampus, we extracted the local theta phase from the microwires by running an ICA over the bundle of eight microwires and selecting the component that had the highest theta (4-8 Hz) component. This peak component was then used to extract the instantaneous phase value 21 ms before stimulation onset by performing a Hilbert transform.

We performed a circular to linear correlation across trials between the phase values extracted from the microwires and the peak amplitude in the macro channels in the post stimulation time-window to determine whether there was any relationship between these two variables. These correlations were z-scored by subtracting the mean value for that micro-maco channel pair and dividing it by its corresponding standard deviation. We then checked whether this value had a chance of p<0.05 to occur given the cumulative distribution function with a mean of 0 and a standard deviation of 1. The total number of significant channel pairs was then counted and evaluated for significance using a binomial test.

To check whether the results of our analysis could result randomly with different frequency values we performed the exact same analysis with the theta values multiplied and divided by 1.618. We chose this value since this value guarantees that the chosen control frequencies never overlap in phase ^39^.

## Supplementary Materials

**Supplementary Figure 1.**
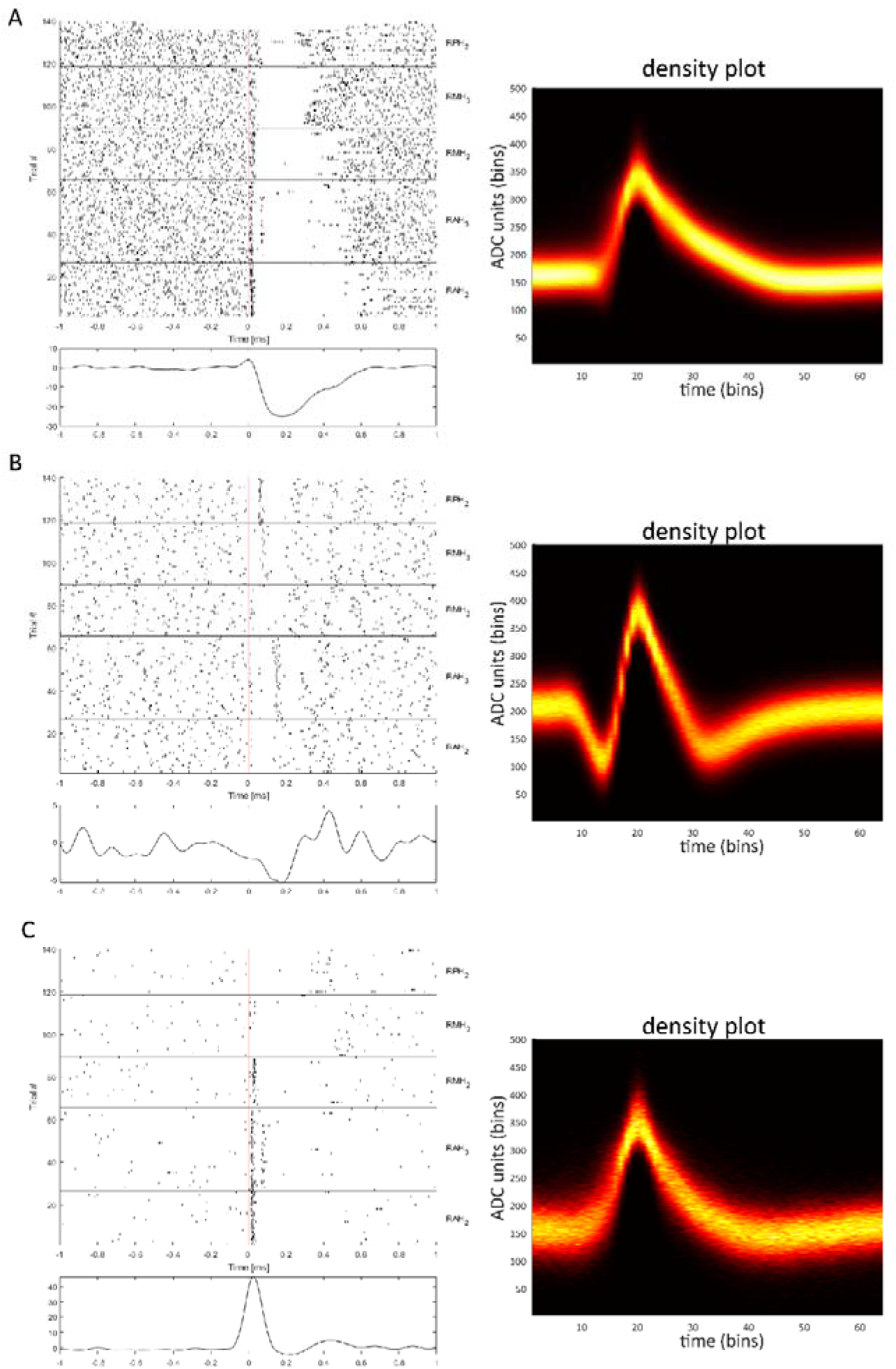
**Example Units:** Examples of three different units exhibiting a significant response to Hippocampal stimulation. Top panels are the raster plots for every single trial separated by the different stimulation conditions (AH,MH,PH, referring to electrodes in the Anterior Hippocampus, Medial Hippocampus, and Posterior Hippocampus respectively). The red line indicates the time point at which the pulse was administered. The lower panel indicates the average GC firing rate over all conditions. The right column contains the density plot of the waveform for the respective unit.

**Supplementary Figure 2.**
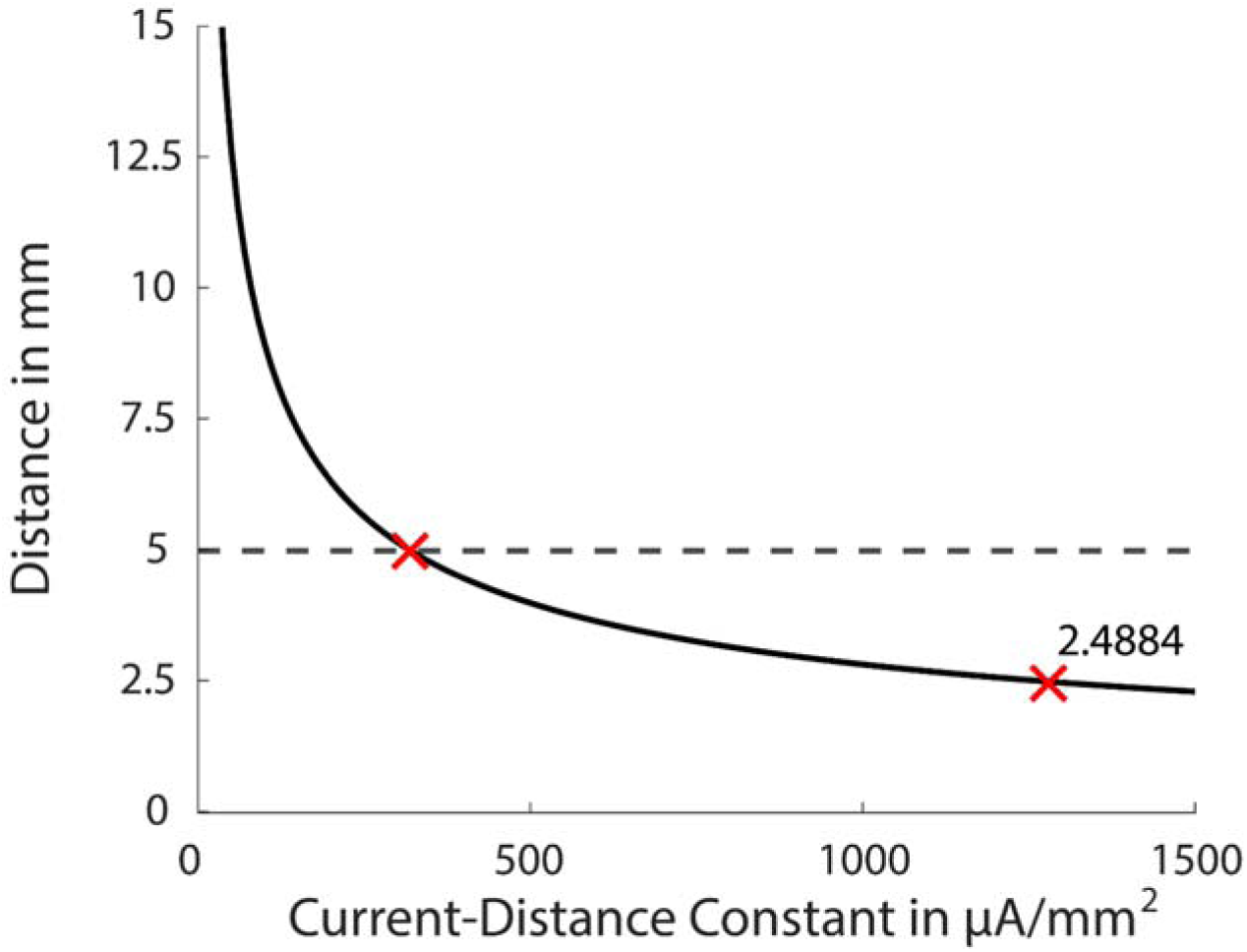
**Electric Spread:** Diagram of the calculated current spread for the administered pulses plotted for each separate current-distance constant. The current distance constant indicates the capacity of neurons to conduct electrical current. This constant has been experimentally assessed in Stoney et al. (1968). This figure was created to see how far off the calculations would have to be for the signal to not be neuronal in nature. The dotted line marks a 5 mm distance as that is the typical distance between the macro channels. The left-most red X highlights the value necessary for crossing the threshold for this distance with a Current-Distance Constant of ∼400 µA/mm². The rightmost red X indicates the averaged current-distance constant determined for cortical neurons in primates. The calculated spread of current was calculated to be ∼2.5 mm using the formula *r = √1/k* ^24,55^. For current spread to exceed a spread of 5 mm the original value would have to be wrong by approximately a factor of 3.

**Supplementary Figure 3.**
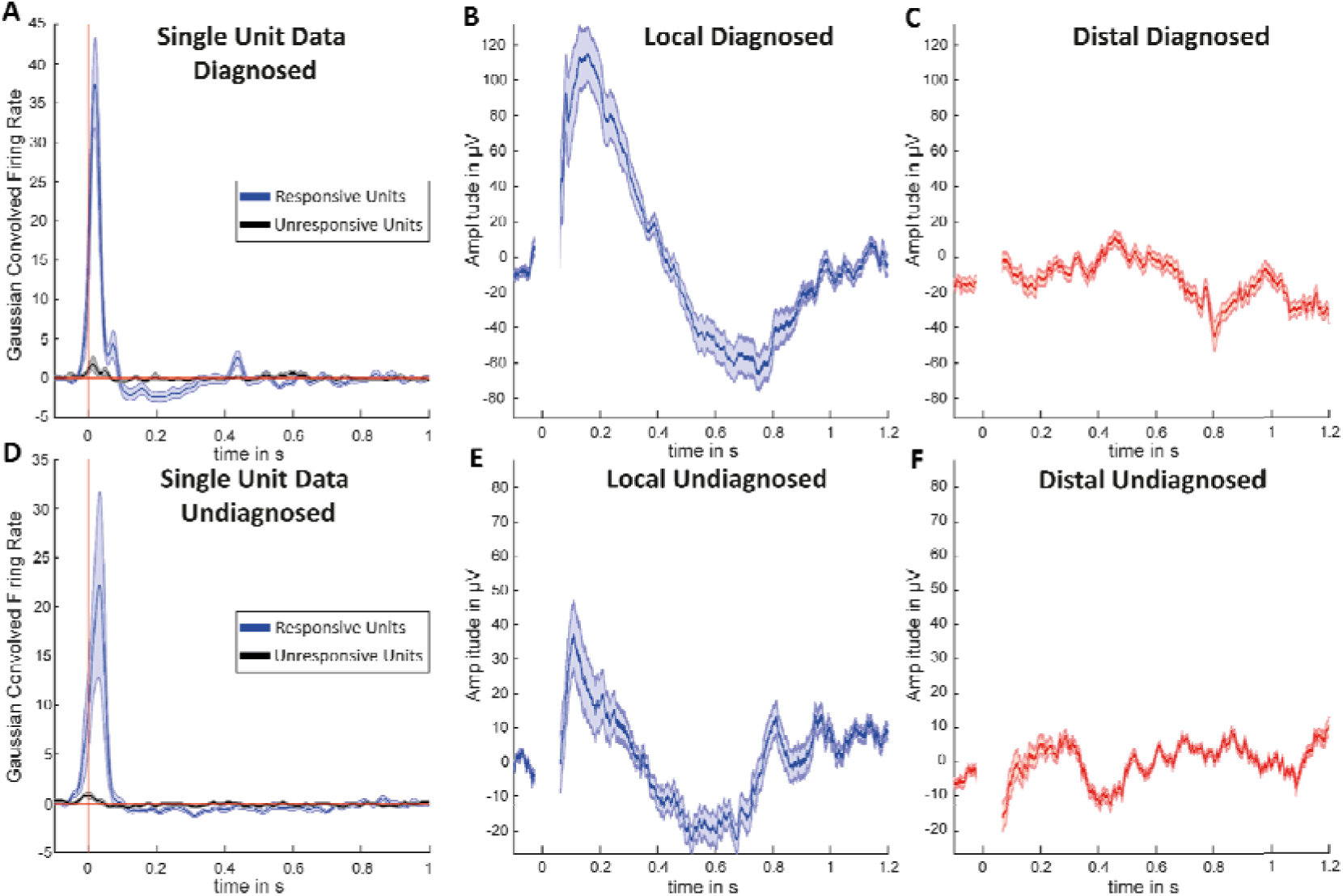
**Controlling for Pathology**: Figure depicting the hippocampal responses to DES. The responses are divided based on whether the tissue in which the electrodes were implanted was diagnosed as epileptogenic (A-C) or not (D-F). Certain sites were diagnosed as epileptogenic if the source of epilepsy was clinically determined to originate in the hippocampus, or adjacent cortical tissue. Single-unit responses show responsive (blue) and unresponsive (black) units in both diagnosed (A; N = 42) and undiagnosed (D; N = 94) regions. Both conditions preserved the characteristic pattern of initial excitation followed by a silent period, confirmed by cluster-based permutation tests (diagnosed: P < 0.007, cluster time = 0.166-0.285 s; undiagnosed: P < 0.001, cluster time = 0.281-0.425 s). Comparison between diagnosed and undiagnosed conditions revealed no significant clusters. Local field potentials recorded via microwires demonstrate responses to local hippocampal stimulation (B, E) and distal cortical stimulation (C, F) in diagnosed and undiagnosed tissue. Cluster-based permutation tests revealed no significant differences between diagnosed and undiagnosed conditions for either stimulation type. Time is shown in seconds; amplitude in microvolts (μV) for field potentials and Gaussian convolved firing rate for single units. Shaded areas represent standard error.

## Acknowledgements

This manuscript was supported by the following grants: European Research Council grant no. 647954 (S.H.), European Research Council grant no. 715714 (M.W.) and European Research Council grant no. 101001121 (B.P.S). The funders had no role in study design, data collection and analysis, decision to publish or preparation of the manuscript. We thank all the patients who have participated in our study.

## Notes

### Competing Interest Statement

The authors have declared no competing interest.

